# Dual SMAD inhibition and Wnt inhibition enable efficient and reproducible differentiations of induced pluripotent stem cells into retinal ganglion cells

**DOI:** 10.1101/682666

**Authors:** Venkata R. M. Chavali, Naqi Haider, Sonika Rathi, Vrathasha Vrathasha, Teja Alapati, Jie He, Kamaljot Gill, Roman Nikonov, Thu T. Duong, Devin S. McDougald, Sergei Nikonov, Joan O’ Brien, Jason A. Mills

**Author notes:** Corresponding authors: Venkata R. M. Chavali, Scheie Eye Institute, University of Pennsylvania, 133A Anatomy Chemistry Building, 3620 Hamilton Walk, Philadelphia, Pennsylvania, USA, 19104., and Jason A. Mills, F.M. Kirby Center for Molecular Ophthalmology and Center for Advanced Retinal and Ocular Therapeutics (CAROT), Scheie Eye Institute, University of Pennsylvania, Perelman School of Medicine, 309B Stellar Chance Laboratories, Philadelphia, Pennsylvania, USA, 19104.

## Abstract

Glaucoma is a group of progressive optic neuropathies that share common biological and clinical characteristics including irreversible changes to the optic nerve and visual field loss caused by death of retinal ganglion cells (RGCs). The loss of RGCs manifests as characteristic cupping or optic nerve degeneration, resulting in visual field loss in patients with Glaucoma. Published studies on *in vitro* RGC differentiation from stem cells utilized classical RGC signaling pathways mimicking retinal development *in vivo*. Although many strategies allowed for the generation of RGCs, increased variability between experiments and lower yield hampered the cross comparison between individual lines and between experiments. To address this critical need, we developed a reproducible chemically defined *in vitro* methodology for generating retinal progenitor cell (RPC) populations from iPSCs, that are efficiently directed towards RGC lineage. Using this method, we reproducibly differentiated iPSCs into RGCs with greater than 80% purity, without any genetic modifications. We used small molecules and peptide modulators to inhibit BMP, TGF-β (SMAD), and canonical Wnt pathways that reduced variability between iPSC lines and yielded functional and mature iPSC-RGCs. Using CD90.2 antibody and Magnetic Activated Cell Sorter (MACS) technique, we successfully purified Thy-1 positive RGCs with nearly 95% purity.

## Introduction

Glaucoma encompasses a heterogenous group of optic neuropathies, united by their exhibition of permanent damage to the optic nerve.^1^ This degeneration is due to the death of retinal ganglion cells (RGCs), with subsequent visual field loss.^2^ Primary open-angle glaucoma (POAG), the most common form of glaucoma, is characterized by chronic and progressive optic nerve degeneration and corresponding visual field deficits in the presence of an open and normal iridocorneal chamber angle.^3^ This disease is the leading cause of irreversible blindness worldwide ^4^, with an estimated 11.1 million expected to become blind from POAG by 2020.^5^ Despite the prevalence of POAG, its pathogenesis remains poorly understood. The complexity of glaucoma certainly makes it possible, if not probable, that RGCs become inherently susceptible to this disease process.

RGCs are found in the ganglion cell layer of the retina and serve as the projection neurons of the retina, utilizing long axons to effectively connect the eye to the brain. They transmit both image-forming and non-image-forming visual information, processed by retinal cells such as photoreceptors and bipolar cells, to higher visual centers in the lateral geniculate body through the optic nerve.^6^ Currently, there are many treatments that slow disease, but no precision treatment exists for glaucoma or RGC degeneration. Although shown to be effective in animal models of glaucoma, neuroprotective approaches have not proven practical in human settings.^7,8^ As mature mammalian RGCs are a terminally differentiated lineage, they do not regenerate after succumbing to disease^9^, consequently leading to irreparable blindness. Understandably, there is a great desire for an application devised to rejuvenate or replace injured RGCs. Hence, it is with good reason that translating human stem cell technology to a viable regenerative therapy for use in degenerative conditions has become a research priority.

Human pluripotent stem cells (hPSCs) offer a tantalizing value in medicine - they can be guided to differentiate into practically any type of cell within an organism. Owing to a variety of techniques that are established to derive every key retinal cell lineage, blinding disorders of the retina can be particularly well-modeled with hPSCs.^10,11^ Though the field was previously reliant on embryonic stem cells (ESCs), advances in stem cell research led to the discovery that overexpressing genes associated with “stemness” in somatic cells can reprogram them into induced pluripotent stem cells (iPSCs).^12,13^ The advent of iPSC manipulation has opened up opportunities for implementing this technology to cure diseases stemming from retinal degenerations.^14^ Taking hPSCs and differentiating them toward RGC lineage commitment may generate enough healthy cells to compensate for the degenerating RGCs in glaucoma. As a result, stem-cell based therapy holds promise as a method to restore vision in conditions of retinal cell loss; however, success of these treatment strategies hinges on de novo synthesis of RGCs with stable phenotypes from hPSCs.^15,16^

Several methods for differentiating ESCs and iPSCs into RGCs have been reported. Of the retinal lineages, RGC differentiation is perhaps the most complicated, historically involving threedimensional (3D) culture of hPSCs.^17,18^ This strategy enables the primordial eye to self-organize into 3D cell aggregates such as embryoid bodies (EBs), followed by optic vesicles (OVs) and later, optic cups, all of which are analogous to structures developed *in vivo*.^18,19^ The retinal progenitor cells (RPCs) eventually mature into RGCs, are characteristically the first neuronal subtype to develop, as well as other retinal cell types.^20,21^ While most of the lineages of the retina do arise in an organized manner, therein lies the problem with the earlier RGC generation protocols: instead of predominantly RGCs, the majority of neural retinal cell types are present.^22–24^ This heterogeneity and inefficient yield in culture restricts selection of specifically RGCs without an enrichment step, even if the 3D platform phase were to precede an adherent culture.^22,25^ To better specify the RGC cell lineage and ideally obviate 3D culture, latter studies modulated RGC signaling pathways, including Hedgehog and Notch, using various ligands and other factors.^24,26^ While these did enhance efficiency, there still resulted a suboptimal RGC yield and an ambiguous picture of which pathways to best modulate, due to the ill-defined addition and administration of components. This shortcoming has been improved upon by administration of small molecules and peptide modulators at appropriate intervals in a calculated manner to precisely modulate cellular signals and pathways such as bone morphogenetic protein (BMP), Wnt, insulin like growth factor-1 (IGF1), and TGF-β.^27^ Canonical Wnt, BMP, and nodal are the three signaling pathways classically inhibited *in vitro* to spur pluripotent stem cells toward the neural retinal fate.^28–30^ Once RPCs form the neural retina, the basic helix-loop-helix transcription factor, ATOH7 plays a central role in RGC commitment and regulation of RGC-specific markers BRN3+ and ISL1.^25,31,32^ Regardless of the protocol, adequate differentiation relies on emulating the pathways present in generating these cell lineages *in vivo*.

Developing a therapeutic approach employing stem cell technology requires an efficient, reproducible, and safe differentiation protocol for RGCs. To this end, we detail a chemically defined *in vitro* technique for cultivating RPC populations from iPSCs that are then directed toward the RGC lineage. True to their *in vivo* development, we directed unaffected control lines of iPSCs to differentiate in a stepwise manner using a temporal induction method centered on small molecule and peptide modulator treatment to inhibit BMP and TGF-β (SMAD), and canonical Wnt, yielding a robust population of iPSC-RGCs after Day 30. Analysis at the molecular and physiologic levels using flow cytometry, immunolabeling, gene expression, and electrophysiology yielded results in accordance with the RGC lineage and allowed specific subtype identification.^24,26,33^

## Methods

### Human iPSC culture

Undifferentiated iPS cells, Control 1 (CHOPWT8), Control 2 (CHOPWT9), and Control 3 (CHOPWT10) were derived and characterized as previously published showing a complete analysis of iPSC characteristics.^34–36^ All human sample collection protocols were approved by the University of Pennsylvania and Children’s Hospital of Philadelphia Human Subjects Research Institutional Review Board following Declaration of Helsinki. All methods were performed in accordance with the relevant research guidelines and regulations of University of Pennsylvania. Written informed consent was obtained from all human cell donors. The iPSC cells were maintained in iPSC medium (Dulbecco’s modified essential medium/ Ham’s F12 nutrient media; DMEM/F12 (50:50; Corning)) containing 1% Glutamax, 1% penicillin/streptomycin (PS), 15% Knockout serum replacement (KSR), 1% non-essential amino acids (NEAA), 0.1mM β-Mercaptoethanol (2-ME) (Life Technologies, CA), and 5 ng/mL of basic fibroblast growth factor (bFGF; R&D Systems) on 0.1% gelatin coated dishes with irradiated mouse embryonic fibroblast (iMEFs).

### Retinal Progenitor Cell generation and conditions

iPSCs were cultured on 0.1% gelatinized plates containing iMEFs in 37°C 5% O_2_ and 5% CO_2_ conditions. Cells were maintained until approximately 75% confluence, then feeder cells were depleted and approximately 1.5 x 10^6^ iPSCs were seeded in one well of 6-well tissue culture dish (Corning) coated with 1:100 diluted growth factor reduced Matrigel. iPSCs were maintained in iPSC: MEF-conditioned medium (80:20) + 20ng/mL of bFGF and 5ng/ml of stable bFGF. The MEF-conditioned media was prepared by plating iMEFs onto 0.1% gelatin at a density of 20,000 cells/cm^2^ in iPSC media. Two days post-plating, media was collected, filtered, and either used directly or cryopreserved for later use. iPSCs were maintained in 37°C at 5% O_2_ and 5% CO_2_ until reaching 100% confluence then transferred to 37°C, 5% CO_2_ overnight prior to induction. On day 0, iPSC: MEF-conditioned media was changed to RPC induction media: DMEM/F12 (50:50), 1% P/S, 1% Glut, 1% NEAA, 0.1mM 2-ME, 2% B27 supplement (w/o vitamin A), 1% N2 supplement, containing 2μM XAV939 (X) (Wnt inhibitor; SelleckChem), 10uM SB431542 (SB) (TGF-β inhibitor; SelleckChem), 100nM LDN193189 (L) (BMP inhibitor; Stemgent), 10mM nicotinamide (Sigma-Aldrich), and 10ng/mL IGF1 (R&D Systems). Cultures were fed daily for 4 days. On day 4, culture media was exchanged with RPC induction media containing: 2μM XAV939, 10uM SB431542, 100nM LDN193189, 10ng/mL IGF1, and 10ng/mL bFGF. Media was changed daily for 12 days. On day 16 of differentiation, seven conditions-1) L, 2) X, 3) 1.5μM CHIR99021 (CHIR) (Wnt agonist; SelleckChem), 4) LX, 5) LSB, 6) LSB-CHIR and 7) LSBX were evaluated for RPC and RGC markers on day 23. Cultures from the following four conditions-1) LSBX, 2) LSB, 3) LX, and 4) 1.5μM CHIR99021 (CHIR) (Wnt agonist; SelleckChem) were further evaluated at 35. An additional five conditions-1) L only, 2) X only, 3) LSB, 4) LSBX, and 5) LSB-CHIR were evaluated for RPC markers on day 23 to evaluate for transcript expression differences across different conditions in RPCs. Cultures were treated for 6 days before transcript analysis.

### Retinal ganglion cell (RGC) differentiation

Prior to RGC differentiation, medium was changed daily using RGC induction media containing SHH (250ng/mL; R&D systems) or SAG (100nM; Tocris), SHH was replaced with SAG after establishing equal competence, FGF8 (100ng/mL, R&D systems) for two days. For RGC differentiation (see Figure 1D and 2B for kinetics), on day 24, cells were manually crossed into small clusters with Leibovitz’s medium containing 34uM Glucose using the crosshatching technique as previously described (Espuny-Camacho et al., 2013), then replated at a density of 1.0 x 10^5^ cells/well of 6-well plate coated with 1:100 diluted growth factor reduced Matrigel in RGCs induction media containing: Follistatin 300 (100ng/ml), cyclopamine(0.5uM, Tocris), DAPT(3uM, Tocris), and 4.2μM Rock inhibitor (Y27632). Media was changed 24h post-plating with RGC induction media containing: Follistatin 300 (100ng/ml) and DAPT(3uM), daily for two days. From day 27, media was changed to RGC induction media containing: DAPT (3uM, Tocris), 10μM Rock inhibitor (Y27632), forskolin (5uM; SelleckChem), cAMP (400uM; Sigma-Aldrich), BDNF (40ng/mL; R&D systems), NT4 (5ng/mL; R&D systems), and CNTF (10ng/mL; R&D systems). Media was changed every 2-3 days until day 36. After maturation (D36), medium was exchanged every 3-4 days with RGC induction media containing: DAPT (3uM, Tocris), 10μM Rock inhibitor (Y27632). Detailed step-wise methodology of iPSC-RGC generation, reagents and concentrations used in our study are provided as supplementary information.

**Figure 1:**
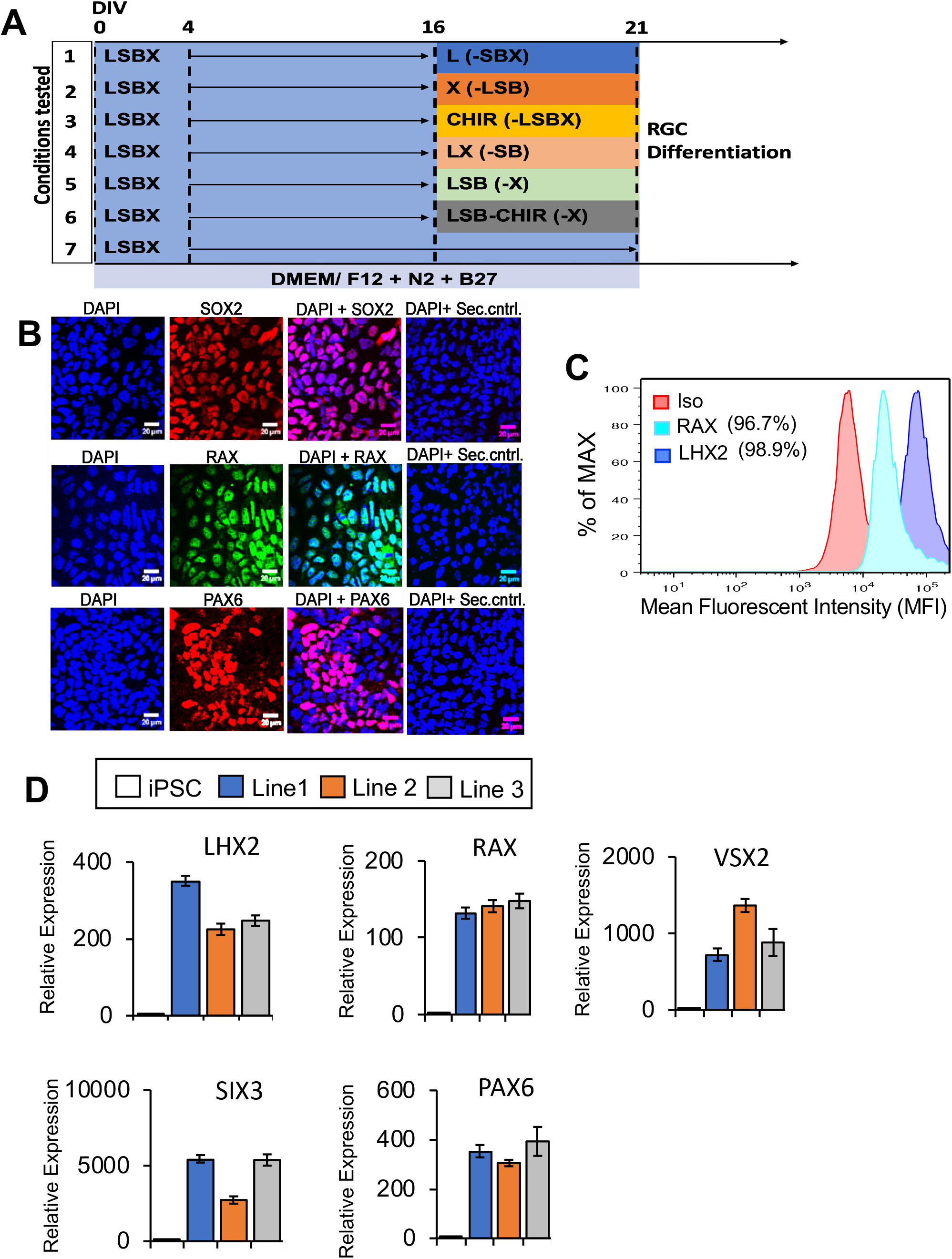
Differentiation of iPSCs into neural retina and retinal ganglion cells (RGCs). A) Schematic showing iPSCs differentiation in neural retinal differentiation media (DMEM/F12 + 2% B27 + 1% N2 + 10 ng/mL IGF1), including inhibitors of BMP, WNT and TGF-β for 15 days, bFGF2 was included in media from day 4-14. At 16 days *in vitro*, cells were treated with RGC induction and maturation media. Differentiation using seven different conditions, L, X, LX, LSB, CHIR, LSB and LSBX are shown. B) Characterization of RPCs using immunocytochemistry with SOX2, RAX and PAX6 along with secondary only control antibodies is shown. C) FACS analysis demonstrated that 96.7% of RPC cells were positive for RAX, and 98.9% of cells were positive for LHX2, when compared to isotype controls. To normalize for flow cytometry cell counts data is represented as %Max. D) Gene expression profiles showing RPC markers in three different iPSC lines, showing a high enrichment. Mean ± SEM represented from at least 3 independent experiments per individual.

**Figure 2:**
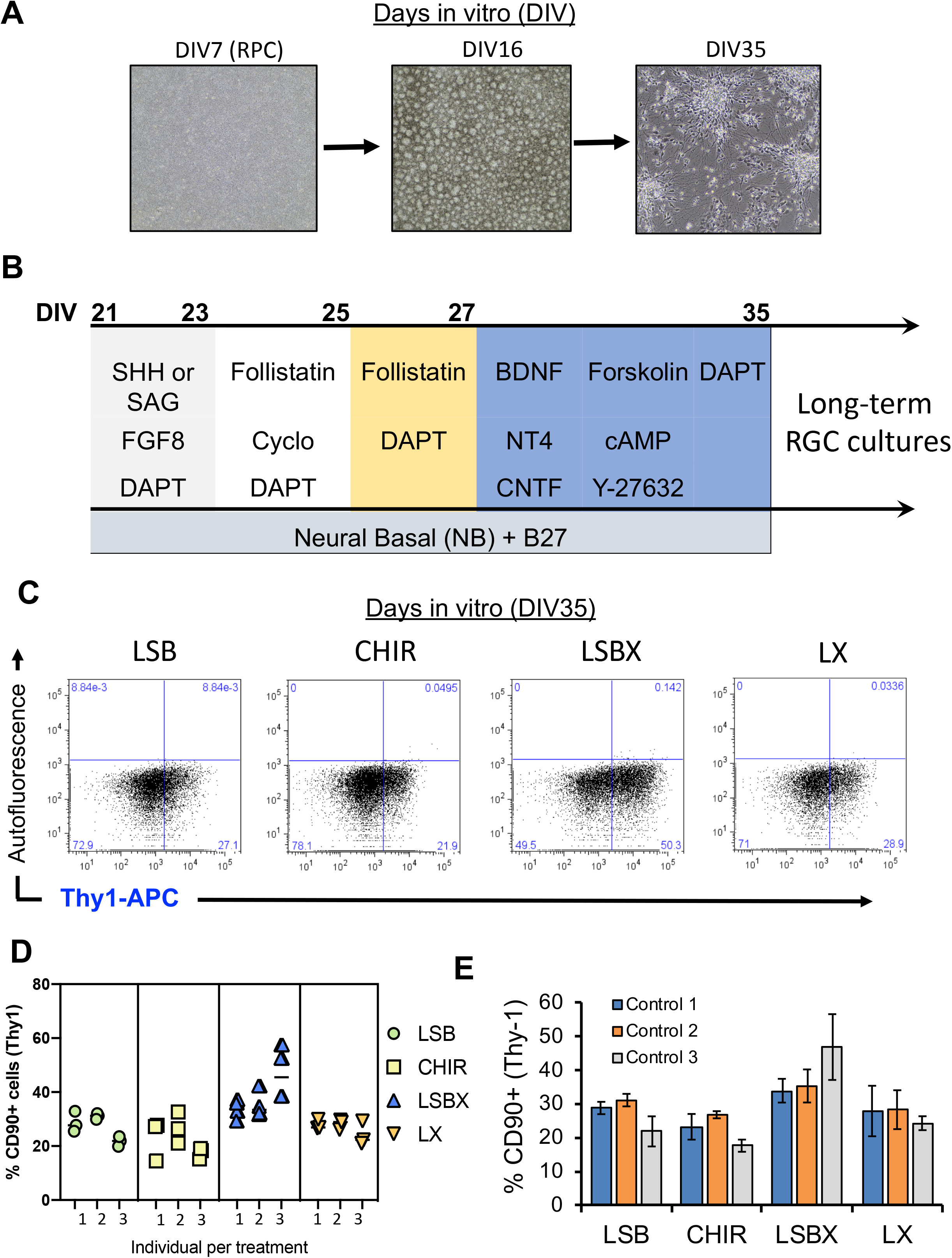
Efficient generation of iPSC-RGCs require inhibition of TGF-β, Wnt, and BMP signaling. A) Phase contract images of different stages of iPSC-RGC differentiation at DIV7, DIV16 and DIV35 is shown. B) Differentiation of retinal progenitor cells (RPCs) to retinal ganglion cells (RGCs) using defined inhibition or activation of developmental pathways, C) Percentage of Thy-1 (CD90) positive iPSC-RGCs in our studies in different differentiation conditions, D) Proportions of CD90 positive RGC cells across four differentiation conditions is shown. Each symbol represents an individual experiment. E) Percentage of CD90 positive cells obtained across three control iPSC lines across each differentiation condition. Mean ± SEM represented from at least 3 independent experiments per individual.

### Flow cytometry analysis of RGCs

iPSC-RGC cultures were lifted using TrypLE (Invitrogen) and collected by centrifugation at 1600 rpm for 5 minutes at 4 °C. The pelleted RGCs were resuspended in 1X HBSS and run through a 100 μm filter to ensure single cell suspension. A portion of cells were incubated with various antibodies anti-CD90-PE-Cy7 (Thy1), anti-Brn-3b Alexa Fluor 647, and anti-RBPMS Alexa Fluor 647 in 1% PBS supplemented with 0.5% bovine serum albumin plus 0.1% Na-Azide (FACS buffer) and incubated at room temperature (RT) for 40 minutes on ice at 4°C as previously described.^41^ We analyzed stained cells using AccuriC6 and data analyzed using FlowJo Version 10.0.8 software (TreeStar).

### Immunohistochemistry

Immunohistochemical analysis was performed as previously published (Duong et al. 2018). For immunolocalization of antigens, RGCs plated on 14mm coverslips were fixed in 4% paraformaldehyde followed by permeabilization using 0.1 % Triton X-100 for 5 minutes. Cells were incubated for 1 hour at RT with a blocking buffer (1 % BSA, 5 % normal goat serum, 0.1 % Triton X-100 in 1 % PBS). RGCs were incubated with primary antibodies diluted in 1 % BSA, 0.1 % Triton X-100 in 1 % PBS buffer for overnight at 4 °C. After washing thrice with 1X PBS, cells were incubated with secondary antibodies diluted in 1 % BSA, 0.1 % Triton X-100 in 1 % PBS buffer for 1 hour at RT. The coverslips were then mounted on slides using prolong Gold mounting medium with DAPI (4’,6-diamidino-2-phenylindole) (Invitrogen). RGCs immunohistochemistry was performed as described earlier ^36–39^ using SOX2, PAX6, RAX, Tuj1, Thy1/CD90, MAP2, GFAP, CRALBP, Brn-3a, and RBPMS antibodies with dilution as provided in Supplementary Table 1. Slides were observed under an Olympus FV1000 Confocal microscope and images were captured with the use of appropriate filters and lasers.

### Voltage recordings

The circular cover glass coated with 1:100 growth factor reduced matrigel containing adherent cells was transferred to the recording chamber filled with Neurobasal media supplemented with 2% B27 without vitamin-A and 1% Glutamax. The chamber was placed on the microscope stage (Olympus BX-61 microscope) and perfused with oxygenated Ames’s solution (Sigma-Aldrich) resulting in gradual replacement of the growth medium with the Ames medium. The temperature of the solution inside the chamber was gradually increased to 37°C prior to imaging and recording session using Warner Instruments TC 344B. Confocal images of the cells were acquired with Olympus Fluoview 1000 MPE system and cells demonstrating strong GFP expression were selected for patch-clamp recording. Whole cell configuration was achieved in the voltage-clamp mode at −60 mV holding potential. After achieving whole-cell configuration, cells were either maintained in the current clamp mode at zero holding current and depolarized with calibrated current steps, or in the voltage-clamp mode at −60 mV holding potential from which they were depolarized with voltage steps. Warner Instruments PC 505B amplifier and Molecular Devices Digidata 1440 digitizer under the control of the Clampex software were used for recording of the membrane voltage and current. Patch pipettes (1.2/0.69 mm) were filled with (in mM) 110 K-gluconate, 12 NaCl, 10 HEPES, 1 EGTA solution. Pipettes were mounted on Sutter MPC-200 micromanipulators; MTI CCD 72 camera system was used to provide video control over pipette and cell positioning in the chamber.

### Quantitative Real-time PCR

Total RNA was extracted using Purelink RNA isolation kit (invitrogen). For each sample, 1μg of total RNA was reverse transcribed using the SuperScript III first stand cDNA synthesis kit with random primers (ThermoFisher). Amplified cDNA was used to quantify Taqman probes (target-FAM and housekeeping-TET) provided in Supplementary Table 4 and TaqMan Fast advanced Master Mix (ThermoFisher) on a 7900 Fast Real-time PCR system (Applied Biosystems). All results were normalized to a *B2M* housekeeping control and were from 3 technical replicates of 3 independent biological samples for each time-point and experimental condition.

### Magnetic Activated Cell Sorting (MACS) to purify CD90+ve RGCs

RGC cells were lifted using TrypLE (Invitrogen), pelleted by centrifugation at 350g for 5 minutes, and total cell number was determined. Cell pellet was resuspended in 90μL buffer (1X PBS pH 7.2, 0.5 % BSA, and 2mM EDTA) and 10μL of CD90.2 microbeads (catalog # 130-121-278, Miltenyi Biotec) per 10^7^ total cells. Cell suspension was mixed well and incubated at RT for 15 min in a tube rotator. In the meantime, MS column was placed onto a MACS separator and the column was prepped. Following the 15 min incubation, the cell suspension was applied onto the column. Flow-through from the column represented the unlabeled or CD90.2^-^ cell fraction. The column was washed with appropriate volume of buffer for at least twice. The column was then removed from the separator and placed on a suitable collection tube. Appropriate volume of buffer was added to the column and magnetically labeled CD90.2^+^ cells were immediately flushed out by firmly pushing the plunger into the column. The cells were plated using RGCs induction media containing 3 μM DAPT and 10 μM ROCK inhibitor.

### Statistical Analysis

Quantitative data were obtained from three independent experiments per cell line in triplicate. Statistical analysis was performed with Student T-test in Prism. \**p*-values of <0.05 were considered statistically significant.

## RESULTS

### Combined inhibition of Wnt, BMP, TGF-β and nicotinamide contributes to efficient generation of early RPCs

The hiPSC retinal differentiation methods can differ between 2D and 3D culture conditions, and their use of proteins and small molecules to recapitulate processes responsible for vertebrate retina development.^26,40^ Previously, the generation of RGCs from hPSCs required that cells be maintained on matrigel as an adherent support to culture cells obtained from 3D cell aggregates and several additive factors like Noggin, DKK-1, IGF-1, bFGF2, and proneural supplements.^41,42,43^ Teotia *et al*, successfully used a modified version of the “Lamba protocol” by differentiating hiPSCs initially into neural rosettes, manual picking, and then induction to functional RGCs, by recapitulating retinal developmental pathways using chemical regulators.^26,41^ These conditions require mid-differentiation enrichment steps that involves the establishment of retinal rosettes followed by manual isolation and maturation to generate retinal subtypes. CRISPR engineered iPSC lines containing fluorescent RFP reporter tag in the *Brn3B* locus, greatly assisted in evaluation of pathways necessary for RGC differentiation and characterization.^44^ This methodology provided a protocol which utilized a monolayer cultures with defined factor supplementations; however, the evaluation were only performed using human embryonic stem cells (hESCs) and resulted in proportions of RGCs between 20-30% of the overall retinal differentiation. A major challenge in the regenerative medicine and disease modeling field are the reproducibility between experiments, and variation between individual to individual. Therefore, we set out to develop and characterize a modified two-stage protocol that differentiates hiPSCs into an enriched population of retinal progenitor cell (RPC) cultures followed by targeted differentiation to RGCs that is reproducible, efficient, and requires minimal personnel interpretation in RGCs generation and maintenance.^26^ To accomplish this, hiPSCs were grown to confluence and subsequently treated with a RPC induction media containing: DMEM/F12 plus N2, B27, XAV939 (WNT inhibitor), SB431542 (TGF-β inhibitor), LDN193189, (BMP inhibitor), nicotinamide, and IGF1 for 4 days (Fig. 1A). The inhibition of Wnt and BMP signaling has been documented to enhance the expression of eye field transcription factors (EFTFs) during retinal differentiations of hPSC.^27^ We observed that addition of TGF-β inhibition induced greater EFTFs expression during early retinal differentiation. Nicotinamide was added to the differentiation media (D0-D3) to promote the expression of early eye field markers LHX2 and RAX, as previously published.^45^ Nicotinamide has been shown to promote cell expansion and adaptation to a radial/rosette morphology.^46^ Differentiation factors such as IGF-1 and bFGF2 aid in the specification of eye field identity to differentiating retinal progenitors.^27^ From Day 4-21, nicotinamide was removed and bFGF was added to RPC induction media. Analysis at day 7 showed an uniform population of SOX2, RAX and PAX6 positive cells (Fig. 1A). The expression of early retinal progenitor markers, LHX2 and RAX, were identified in over 95% of day 7 cultures (Fig. 1B) indicating an efficient and robust generation of RPCs. Quantification of EFTFs, *LHX2*, *RAX*, *PAX6*, *SIX6*, *SIX3*, and *VSX2*, showed significant expression relative to pluripotent stem cells in all three individual cell lines (Fig. 1C).

The efficient induction of retinal progenitors is largely dependent on the inhibition of pathways responsible for patterning of diencephalon; however, it is unclear if modification of these inhibitory pathways play a role on retinal ganglion cell generation. Here, we tested whether pharmacological inhibition of Wnt, TGF-β, and BMP can improve RGC induction and subsequently enable the controlled generation of mature iPSC-RGC neurons. We evaluated seven different treatment conditions to determine pathways that enhance or restrict RPC generation from iPSCs. The following 7 conditions were tested from d16-d21: 1) BMP (L), 2) Wnt inhibition (X), 3) CHIR (Wnt agonist), 4) LX, 5) BMP and TGF-β (LSB), 6) LSB-CHIR and LSBX (Schematic in Fig. 1A).

### Extended exposures of BMP, Wnt, and TGF-β inhibition leads to efficient generation of RGC neurons

RGC differentiation proceeded through two phases: 1) uniform RPC cultures (up to D21), and 2) late stage RGC differentiation (D22-36) (Fig 2A,B). At day 22-23, cells were treated with RGC induction conditions that included the activation of sonic hedgehog, by augmenting with SHH (250 ng/mL). Comparative studies in our lab showed the use of SAG (100nM, smoothened agonist) could replace SHH with no difference in RGC induction changes. Fibroblast growth factor 8 (FGF8) and notch inhibitor (DAPT) were added along with SHH or SAG. During iPSC generation, SHH pathway signaling was shown to enhance the specification of RPCs to RGCs during this 48h time frame (supplementary information)^47^. On day 24, cultures were expanded using a cross-hatching technique known separate cells into small clusters and SHH, TGF-β, and notch signaling were inhibited to promote RGC maturation using cyclopamine, follistatin, and DAPT, cyclopamine was removed for the following 2 days. To improve survival of RGCs during the expansion,^48^ cAMP, BDNF, neurotrophin 4 (NT4) and CNTF, were added to the media to promote the development and survival of cells with neuronal fate.^49^ Forskolin and Y-27632 were included as compounds shown to promote RGC neurite growth, survival and lineage commitment.^40^ These conditions were established by combining previously described conditions used by Sluch *et al*., 2015 and Teotia *et al*., 2017 for hiPSC-RGCs. After 35 days *in vitro*, cultures were disassociated and flow cytometric analysis were performed on single cell suspension obtained from three independent iPSC lines, and data showed that highest proportions of RGCs were generated from the LSBX conditions, with a range between 40-50% Thy-1 (CD90) (Fig. 2C, E), 82-84% Brn-3b, and 11-12% RBPMS(Supplementary Fig.1). The LX, CHIR, and LSB conditions resulted in 26.7% (±3.2), 22.5% (±4.7), and 27.2% Thy-1 positive cells (±6.2), respectively (Fig. 2C). We observed consistent results across individuals and experiments (Fig. 2D).

Additionally, on Day 35 we further concentrated the number of RGCs in our culture by employing the MACS technique and CD90.2 microbeads. Since RPCs are multipotent cells, they have the potential to differentiate into neuronal cell types and one glial cell type called the Müller glial cells ^61,62^. Therefore, we utilized the CD90.2 microbeads for the positive selection of RGCs expressing Thy1 cell surface marker and the removal of Müller glial cells from our culture. Following the purification of RGCs using MACS, the cells were analyzed using immunocytochemistry by detecting for the presence of Müller glial cells, astrocytes, and RGCs using CRALBP, GFAP, MAP2 TUJ1, RBPMS and Brn-3a markers, respectively (Fig. 3). Our results show that roughly 95% of the cells in our culture were positive for Brn-3a with the presence of extended synaptic connections between RGCs. Whereas, about 5% of the cells were positive for astrocytes and we detected no presence of Müller glial cells in our system (Fig. 3). However, the presence of astrocytes is significant since they are essential for the functional activity of the neuronal cell types by ensuring proper synaptic maturation and signaling^63^.

**Figure 3:**
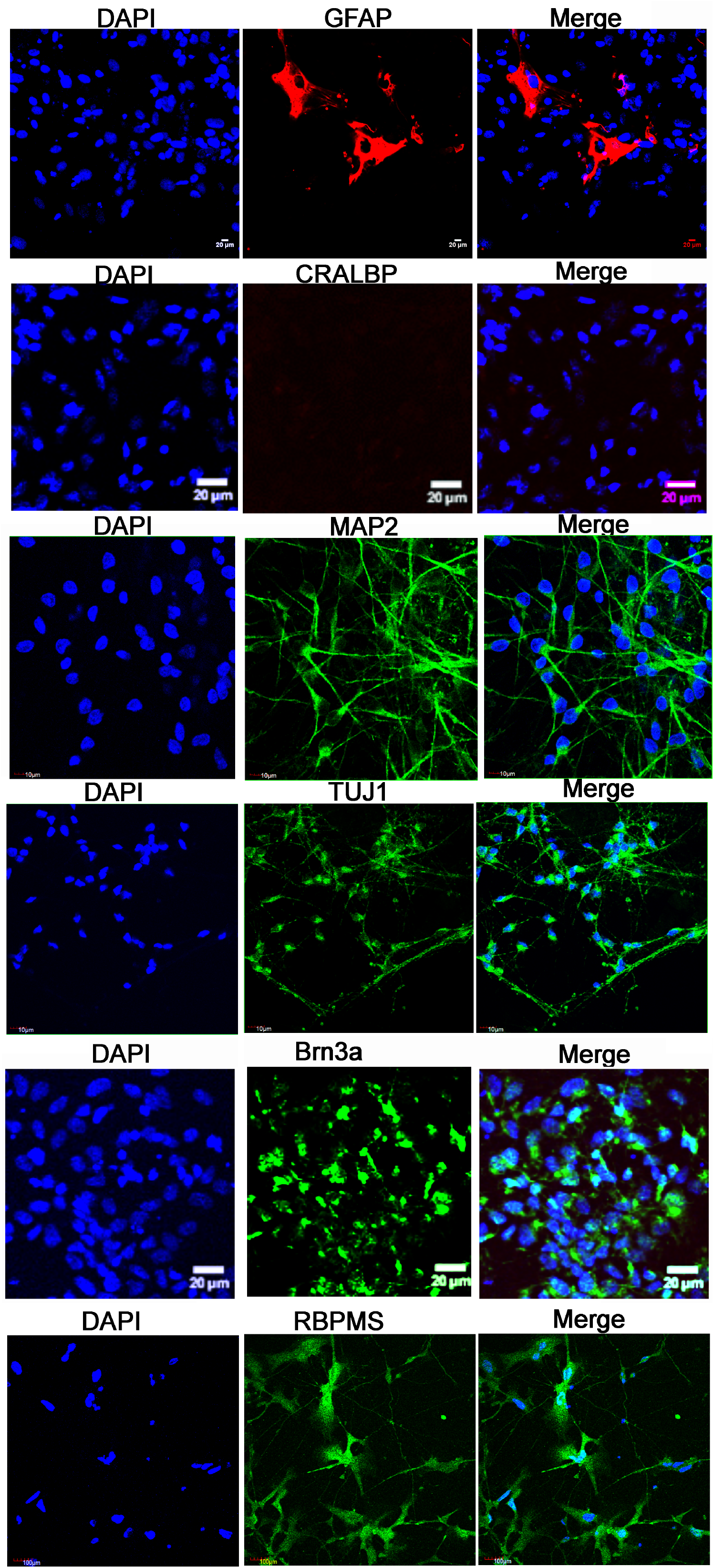
Charecterization of iPSC-RGCs after MACS sorting using ICC. The iPSC-RGCs stained positively to RGC markers, Brn3a, TUJ1, MAP2, RBPMS. Postive staining with GFAP showed <5% astrocytes like cells. No cells stained positive with CRALBP.

For long-term cultures, we initially used RGC induction supplemented with 1% N2 and 1% B27 for the maintenance of RGC cultures after DIV35; however, this resulted in significant overgrowth of proliferative late RPCs. The removal of N2 supplementation reduced the number of dividing cells in our cultures, providing a culture with predominantly RGCs. We have also cultured RGCs in 1% CultureOne supplement (Life Technologies) in the RGC maturation media, which reduced the number of non-RGC cell growth in day 45 (only in two weeks, from Day 27-40) cultures and beyond. After Day 35 in culture, the iPSC derived RGCs produce appropriate morphological and physiological features of mature RGCs.

### Gene expression profiles during differentiation of iPSC to RPCs and RGCs

During eye formation in vertebrates, cell intrinsic signals, extrinsic signals and/or transcription factors control the differentiation and fate determination of retinal cells. We evaluated the gene expression profile of three EFTs, *RAX*, *PAX6*, and *SIX3* that play a role in the anterior neural plate (Fig. 4). The expression of Rx (encoded by *RAX* gene) was maximum in the RPC inhibited by BMP and Wnt inhibition when compared to the other conditions at DIV23 (Fig. 4, Supplementary Table 5). The *RAX* expression at DIV35 RGCs was minimal suggesting a commitment to a more differentiated retinal fate, a conseqnuence of retinal progenitor cell (RPC) expansion. PAX6 is expressed in the cornea, lens, ciliary body, and retina through development and plays a role in determining their cell fate. The *PAX6* transcript expression was observed in all experimental conditions in RPCs and RGCs; however, predominant expression of *PAX6* is detected at DIV23 and DIV35 in the CHIR condition (Fig. 4), which stimulates the canonical Wnt signaling. Our results indicate that prolonged stimulation of RPCs with Wnt restricts their differentiation potential and maintains majority of the cells as multipotent progenitors.

**Figure 4:**
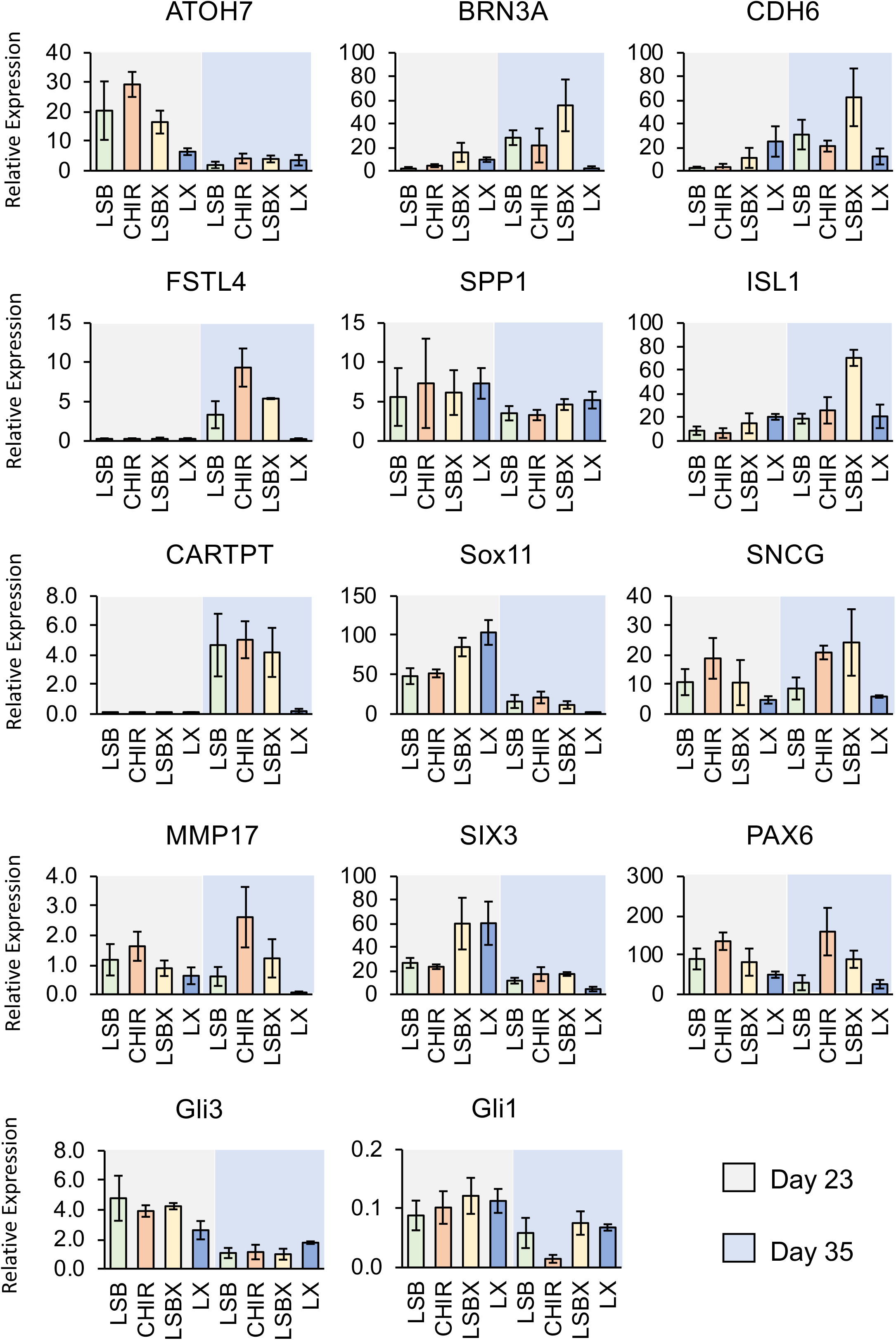
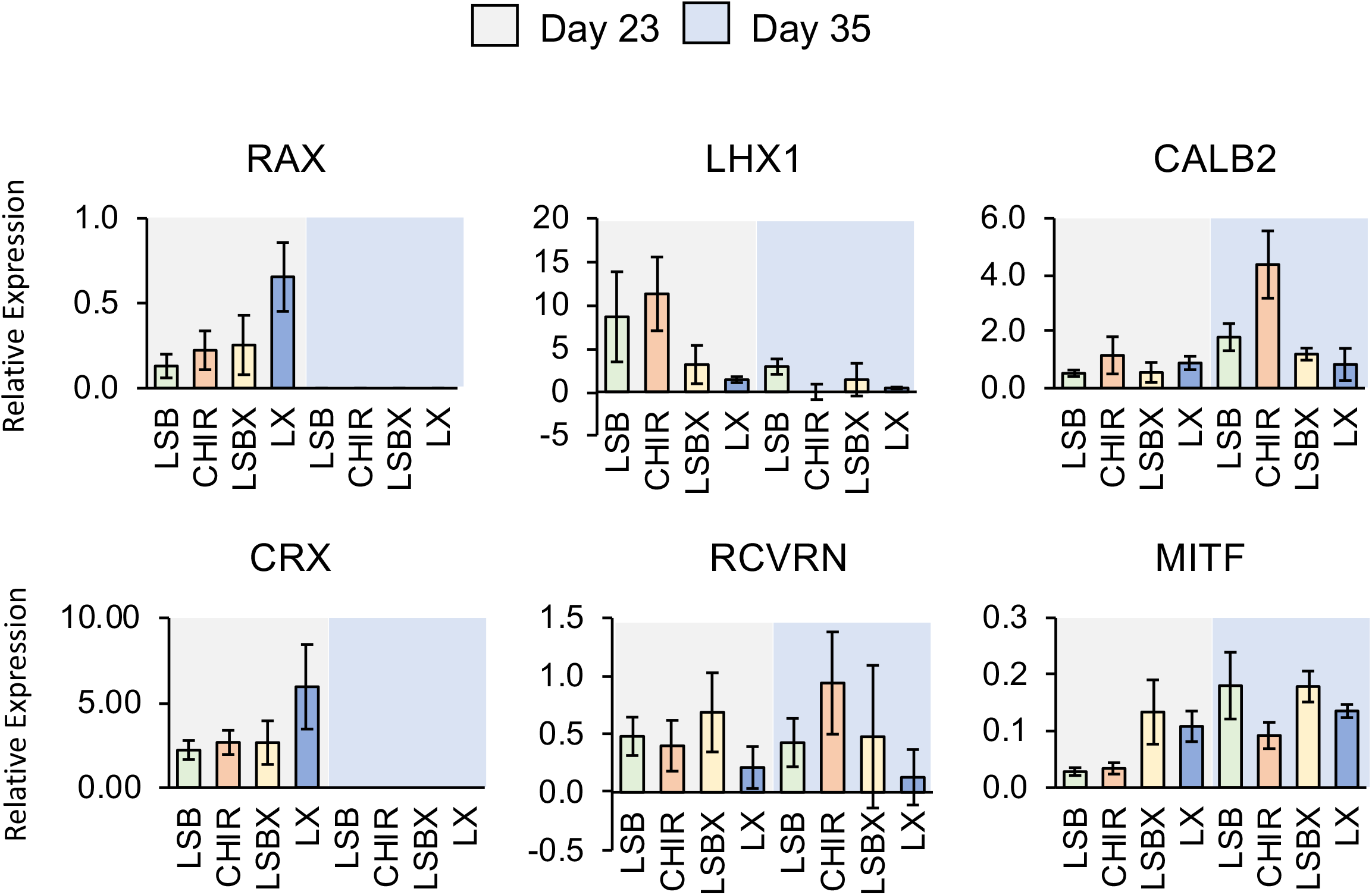
Transcriptional Profiling of iPSC-RPCs. Quantitative-RT-PCR profiles showing relative expression of retinal ganglion cell and retinal subtypes (photoreceptors, amacrine, horizontal, and RPE) at Day 23 and Day 35 stages of development from iPSCs to RGCs.

Among other gene transcripts analyzed, we observed an increase in *SIX3* expression in both LX and LSBX conditions at DIV23 days, when compared to RPCs stimulated with LSB or CHIR. The expression of *SIX3* decreased in RGCs at DIV35 indicating that neurospecification was reaching completion at this stage.

The *SOX11* transcript is heavily expressed in developing retina during embryonic stages^50^. It is required for the maintenance of hedgehog signaling and is critical for axonal growth, extension and driving adult neurogenesis^51–53^. The expression of *SOX11* significantly increased in LSBX and LX conditions when compared to LSB and CHIR in RPCs at DIV23. The expression of *SOX11* was significantly reduced in RGCs at DIV35 (Fig. 4). GLI3 has a dual function as a transcriptional activator and a repressor of the sonic hedgehog (Shh) pathway. GLI1 is a simple transcriptional activator encoded by a target gene of Shh signaling. We observed increased expression of *GLI3* in RPCs at DIV23 when compared to RGCs at DIV35. Expression of both *GLI3* and *GLI1* genes demonstrated that sonic hedgehog signaling is important for development of RGCs.

Direction selective RGC (DS-RGC) are subtypes of RGCs that respond to motion of light in different directions can be identified by expression of specific molecular markers, such as *CART*, *CDH6*, and *FSTL4*, among other genes.^54–57^ The CDH6 mediates axon-target matching and promotes wiring specificity that does not lead to image formation in the mammalian visual system. Cadherin mediated cell-cell adhesion ensures precise connectivity of neurons in the eye to target nuclei in the brain. Increased expression of *CDH6* was observed in LSBX condition in DIV35 RGCs demonstrating their maturation towards increased specificity for axonal wiring between RGCs. The CARTPT (cocaine- and amphetamine-regulated transcript) is expressed by a major subtype of RGCs, ooDSGCs. In our study, the expression of *CARTPT* was seen in RGCs matured only in LSB, CHIR and LSBX conditions at DIV35 indicating that these conditions develop specific subtypes of mature RGCs that are known to be markers for ON-OFF direction-selective RGCs.^58^ *FSTL4* is a gene expressed in ON DS-RGCs and is colocalized with BRN3B in few RGCs.^43^ We have seen increased expression of *FSTL4* gene transcript in CHIR induced RGCs when compared with other conditions at DIV35 indicating the development of various subtypes of mature RGCs in our culture conditions. We did not observe expression of *CARTPT* or *FSTL4* transcripts in RPCs at DIV23 (Fig. 4).

The RGCs at DIV35 expressed *BRN3A* transcript predominantly in LSBX growth condition when compared to other conditions. Interestingly, *ATOH7* expression was observed early in RPC differentiation at DIV23 and decreased in RGCs by DIV35. We detected low *CRX* expression levels in DIV35 RGC cultures across all culture conditions implying that the RGC differentiation conditions at this stage restrict the photoreceptor precursor cell populations. We observed low expression of *RCVRN* (expressed by photoreceptors) and *MITF* in our cultures at DIV23 and 35 indicating that our cultures are differentiated predominantly towards RGC fate, with minimal retinal pigment epithelium cell identity (Fig. 4). Development of few interneurons in mature RGCs (amacrine or horizontal cells) was also observed, as evidenced by expression of *CALB2* especially in RGCs with Wnt activation at DIV35.

To understand the importance and requirement of BMP, Wnt, TGF-β inhibition in differentiation and maintenance of RPCs, we performed pairwise gene expression analysis of the RPC cultures at DIV23 for the four conditions (L, SB, X and CHIR) in different combinations (Supplemetary Table. 5). We observed increased expression of *RAX*, *PAX6*, *LHX2* and *SOX11* at DIV23 in L and X when compared to LSB, LSBX and LSB-CHIR suggesting that BMP and Wnt inhibition is required for maintaining majority of cells in retinal progenitor state. High expression of *CRX*, and *RCVRN* at DIV23 in L, X and LSB when compared to LSB-CHIR suggests that Wnt signaling may play role in preventing the differentiation of RPCs toward photoreceptors lineage. Decreased expression of *CALB2* at DIV23 in X (Wnt inhibitor) condition when compared to LSB, LSBX and LSB-CHIR; while increased expression at DIV35 under CHIR (Wnt agonist) condition suggests that Wnt signaling is crucial for promoting RPC differentiation to RGC. Increased expression of *GLI1* and *GLI3* transcripts at DIV23 in L (BMP inhibitor) condition when compared to LSB, LSBX and LSB-CHIR suggests that BMP inhibition is important for promoting SHH signalling (Supplemetary Table. 5).

### Functional Analysis of iPSC-RGCs

Confocal imaging of iPSC-RGC cultures at DIV35 show the presence of the GFP-expressing neurons with neurite projections, and an increase in the density and complexity of the projections over the period of one month. The iPSC-derived RGCs exhibited morphological features, with large somas connected by elongated axonal processes. Immunocytochemical analysis shows the expression of transcription factors like BRN3A, BRN3B, SNCG, RNA-binding proteins like RBPMS, CD90/THY1 and other cytoskeletal markers like MAP2 and TUJ1; signifying the characteristic of RGCs and RGC expression (Fig. 5A).

**Figure 5:**
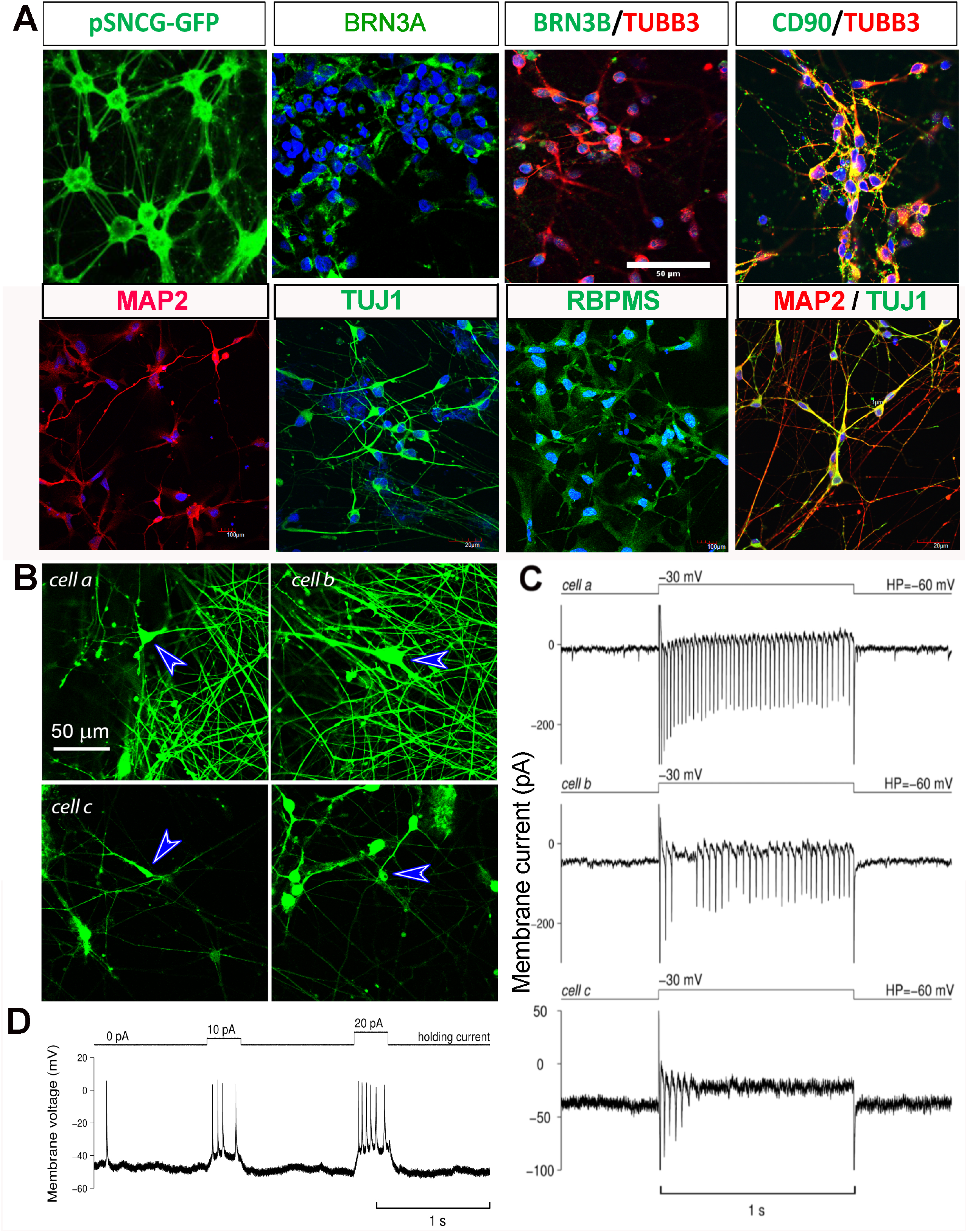
Functional Characteristics of Induced Retinal Ganglion Cells. A) RGCs were transduced with AAV7m8-SNCG-eGFP and immunofluorescence RGC markers (BRN3a, THY1, TUBB3, MAP2, TUJ1, RBPMS). B) Density of RGCs expressing GFP and other axonal processes increases between DIV35 (4B, cell c) and maturation DIV75 (4B, cells a and b). Blue arrows point to cells selected for patch clamp recordings and that fired action potential upon depolarization. C) Action potential firing in response to depolarizing voltage step recorded from cell a and cell b (DIV75) and from cell c (DIV35). D) This trace demonstrates action potential firing in response to the two different depolarizing steps of current recorded from *cell a* from Fig. 4B. We also observe a spontaneous spike firing at the start of the trace. Same scale bar applies to all images.

Two out of the six cells tested demonstrated action potential firing in the current clamp mode in response to the depolarizing currents. The responses of one of these cells (*cell a*) are illustrated in Fig. 5B. Although cells are not expected to fire action potentials under ideal voltage-clamp conditions, we believe that imperfect clamping in distal processes can lead to such firing observed as illustrated in Fig. 5C for the same *cell a* from Fig. 5D. Interestingly, all but one cell (6 out of 6 and 4 out of 5 DIV35 old cells) demonstrated reliable firing when depolarized with voltage steps in the voltage-clamp mode. In Fig.5B, *cell b* illustrates firing of the older cells and *cell c* illustrates firing of the younger cells in response to the depolarizing voltage steps. As expected, later stage iPSC-RGCs cultures (D75) produced higher frequency, sustained firing and generated larger spikes when compared to early born iPSC-RGCs.

Cells firing under current-clamp conditions had resting membrane potentials around −50 mV. In contrast, cells that fired under voltage-clamp but not current-clamp conditions had resting potentials around or above −30 mV. In voltage clamp mode membrane potential was maintained at −60 mV allowing more effective recovery of sodium channels from inactivation after depolarizing steps, thus enabling action potential firing in response to depolarization.

## Discussion

In the current study, we employed a two-step/stage differentiation to induce RGC differentiation from iPSCs by modification of existing methodologies.^26,27,43^ The first stage involved differentiation of iPSCs to RPCs. The RPCs were matured in a stage-specific manner using small molecules and recombinant proteins to modulate SHH pathway, Wnt pathway and Notch signaling to produce abundant RGCs reliability, which stained positive for RGC markers and emulated action potential.

This method is a quick and efficient RGC generation protocol without the need for 3D aggregate formation or manual enrichment to initiate RGC differentiation. In our method, the entire hiPSCs monolayer was differentiated to RPCs using a chemically defined medium in 2D cultures. We employed crosshatching technique to generate clumps of cells that underwent stage-specific differentiation to produce functionally mature RGCs by DIV28. This provided an accelerated timeline considering other published methods to date.^22,26,44,59,60^ Our methodology involved using a chemically defined media standardized by Teotia *et al*, with minor modifications, ^26^ to take advantage of 15-day RGC differentiation timeline and pairing it with our novel RPC generation protocol to produce RGCs. The RPCs generated showed immunoreactivity to RAX and LHX2 in the RPC lines for over 97% of cells in iPSC culture indicating that our protocol committed cells towards RPC lineage. Using our method, we reliably differentiated six normal iPSC lines and multiple clones from those lines (data not shown) to generate mature RGCs that stain positive for Thy1/Tuj1, BRN3A, BRN3B, TUBB3, RBPMS, MAP2 and SNCG markers. Using different chemical conditions, we generated over 80% of pure iPSC-RGC cultures. Using FACS sorting analysis, we quantified the presence of Thy1 positive RGCs (~58%), Brn3b positive (~84%) and RBPMS positive (~12%) iPSCRGCs in our differentiated cultures. Howerver, there is a discrepancy between the FACS quantification and immunocytochemistry (ICC) results for RBPMS positive RGCs in our cultures that could be primarily due to differences in the antibodies used for FACS and ICC and their antigen recognition abilities. Although we are certain about RBPMS antibody used for ICC as it is well characterized in previous studies ^64^. Therefore for FACS analysis we will testing other RBPMS antibodies in future to improve the efficacy of iPSC-RGC detection by this technique.

In addition to the matured RGCs, we also generated fewer other retinal cell types that express pan-retinal markers and appear to be astrocytes, amacrine and/or bipolar cells. Further characterization of these cells is needed to confirm their identity. We did not produce RPE and photoreceptor outer or inner segments in our differentiations.

Here, we also show valuable data representing a heterogenous population of RGCs that can be characterized by cell type specific gene expression. Future work will be to further characterize *in vitro* conditions that will better segregate RGC cell patterning of these subtypes, and to perform single-cell RNA sequencing to confirm cell identity. This will be extremely useful in providing a runway for identification of possibly new surface markers specific to RGC subtypes, which may allow for selection using magnetic bead isolation, fluorescent activated cell sorting or immunopanning of iPSC derived RGCs.

Furthermore, the RGCs differentiated using the LSBX condition exhibited electrophysiological function, with the ability to conduct sodium and potassium through voltage-dependent channels and fire action potentials. When comparing action potentials obtained from RGCs at day 35, day 75 and day 110, we observed that RGCs were able to fire continuously producing larger spikes with higher frequency, as they matured and aged in culture when compared to younger cells. Based on the nature and type of physiological responses, several RGC subtypes like ON-OFF-and alpha-RGCs were observed. Therefore, our validated methodology can reliably harness iPSC technology as a renewable source of RPCs to efficiently produce highly enriched populations of RGCs for *in vitro* studies of glaucoma and potential therapeutic modalities for incurable RGC-related diseases.

## Conclusion

In this study, we present a reproducible and efficient chemically defined *in vitro* methodology to generate unprecedented yields of RPC populations from multiple iPSCs, that are then directed toward the RGC lineage. We differentiated multiple iPSC lines into RGCs in a step-wise manner using small molecules and peptide modulators by inhibiting bone morphogenetic protein, TGF-β, and canonical Wnt pathways. Purified populations of these mature iPSC-RGCs have the potential for *in vitro* studies of glaucoma and for therapeutic purposes for many RGC-related diseases.

## Acknowledgements

Supported in part by the National Eye Institute of the National Institutes of Health under Award Number R21EY028273-01A1 (VRMC), R21EY027936-01A1 (JAM), and 1RO1EY023557-01 (JMO). BrightFocus Foundation grant (VRMC), Research to Prevent Blindness Unrestricted Grant Funds to Scheie Eye Institute (VRMC, JAM, JMO), F.M. Kirby Foundation, and The Paul and Evanina Bell Mackall Foundation Trust for their support The Center for Advanced Retinal and Ocular Therapeutics (CAROT), The Foundation for Fighting Blindness (FFB), Research to Prevent Blindness (RPB), Lisa Dean Moseley Foundation, the National Institute of Health vision core grant P30EY01583-26, National Institutes of Health, Department of Health and Human Services, under eyeGENETM and contract Nos. HHSN260220700001C and HHSN263201200001C.

## Author contributions

VRMC proposed the project and developed the concept with JAM. VRMC, NH, SN, TTD, SR, JAM conceived the method. VRMC, NH, SR, SN, RN, KG, JH, TTD, JAM conducted the experiments. DM generated the SNCG-GFP construct. VRMC, JAM, TTD, SR, NH, and SN performed the data analysis. TA, VRMC and JAM wrote the manuscript. VRMC, JAM, TTD, SR, NH, SN, RN, JMO, JH and TA revised the manuscript. VRMC, JMO and JAM supervised this work. All authors read and approved the final manuscript.

## Additional information

Competing financial interests: The authors declare no competing financial or non-financial interests.

