## Supplementary Information for "Dual SMAD inhibition and Wnt inhibition enable efficient and reproducible differentiations of induced pluripotent stem cells into retinal ganglion cells"

**Supplementary Tables 1-4: Antibodies, small molecules, media reagents and primer sequences used for generation of iPSC-RGCs.**

| **Supplementary Table. 1. Antibodies, their source and concentration used in our study** | | | | | | |
| --- | --- | --- | --- | --- | --- | --- |
| **Protein species** | **Antibody Source** | **Clone/Catalog No** | **Species** | **Concentration/**  **Volume** | **Dilution** | **Application** |
| SOX2 | Cell Signaling Technology | 3579S | Rabbit |  | 1/100 | ICC |
| PAX6 | Biolegend | PRB-278P100 | Rabbit | 2 mg/ml | 1/100 | ICC |
| RAX | Abcam | ab23340 | Rabbit |  | 1/100 | ICC |
| THY1/CD90 | R&D Systems | AF2067 | Sheep | 25μg | 1/100 | ICC, Flow cytometry |
| BRN3b | Santa cruz | sc-514474 | Mouse | 200 μg | 1µg/10^6^ cells | Flow cytometry |
| RBPMS | Novus Biologicals | NBP2-73835AF647 | Mouse | 0.64 mg/ml | 1µL/10^6^ cells | Flow cytometry |
| TUJ1 | Biolegend | MRB-435P | Rabbit | 1 mg/ml | 1/250 | ICC |
| RBPMS | Sigma- Aldrich | ABN1362 | Rabbit | 100 μg | 1/1500 | ICC |
| MAP2 | Sigma | M1406 | Mouse | 2 mg/ml | 1/500 | ICC |
| BRN3a | Santa cruz | sc-8429 | Mouse | 200 µg/ml | 1/100 | ICC |
| BRN3b | Santa cruz | sc-6026 | Goat | 100 µg/ml | 1/100 | ICC |
| TUBB3 | Biolegend | PRB-435P | Rabbit | 1.0 mg/ml | 1/100 | ICC |
| CRALBP | Abcam | ab15051 | Mouse | 1.0 mg/ml | 1/1000 | ICC |
| GFAP | Abcam | ab7260 | Rabbit |  | 1/1000 | ICC |
| SSEA4-APC | Biolegend | MC-813-70 | Mouse | 25μg/ml | 1/100 | Flow cytometry |
| BRN3a | Santa cruz | sc-31984 | Goat | 100μg | 1/100 | ICC (after MACS) |
| MAP2 | Santa cruz | sc-74421 | Mouse | 200 μg | 1/100 | ICC (after MACS) |
| GFAP | Abcam | ab7260 | Rabbit | 50 µl | 1/1000 | ICC (after MACS) |
| Alexa Fluor 594 Goat Anti- Mouse | Invitrogen | A11020 | Goat | 500 μg | 1/500 | ICC |
| Alexa Fluor 488 Chicken anti- Goat | Invitrogen | A21467 | Chicken | 2 mg/mL | 1/500 | ICC |
| Alexa Fluor 488 Goat anti- Rabbit | Invitrogen | A11034 | Goat | 1 mg | 1/500 | ICC |

*ICC- Immunocytochemistry

| **Supplementary Table. 2. Small molecules and protein source information** | | | |
| --- | --- | --- | --- |
| **Small molecules/recombinant protein** | **Company** | **Cat No.** | **Reconstitution** |
| Recombinant Human FGF basic (146 aa) Protein | R & D Systems | 233-FB | 0.1% BSA in PBS |
| Recombinant Human  Sonic Hedgehog/Shh Protein | R & D Systems | 8908-SH | 0.1% BSA in PBS |
| Recombinant Human/Mouse FGF-8b Protein | R & D Systems | 423-F8 | 0.1% BSA in PBS |
| DAPT | Stemgent | 04-0041 | DMSO |
| Recombinant Human Follistatin 300 Protein | R & D Systems | 669-FO | 0.1% BSA in PBS |
| Cyclopamine | R & D Systems | 1623/1 | DMSO |
| Recombinant Human BDNF Protein | R & D Systems | 248-BDB | 0.1% BSA in PBS |
| Forskolin | Selleckchem | S2449 |  |
| Recombinant Human NT-4 Protein | R & D Systems | 268-N4 | 0.1% BSA in PBS |
| Recombinant Human CNTF Protein | R & D Systems | 257-NT | 0.1% BSA in PBS |
| cAMP  N^6^,2′-O-Dibutyryladenosine 3′,5′- cyclic monophosphate sodium salt | Sigma-aldrich | D0627 | DMSO |
| Y-27632 dihydrochloride | R & D Systems | 1254/1 | DMSO |
| LDN 193189 dihydrochloride | R & D Systems | 6053 | DMSO |
| SB 431542 | R & D Systems | 1614 | DMSO |
| CHIR 99021 | Selleckchem | S2924 | DMSO |
| XAV 939 | R & D Systems | 3748 | DMSO |
| Nicotinamide | Sigma-aldrich | N0636 | DMSO |
| IGF | R & D Systems | 291-G1 | 0.1% BSA in PBS |
| CHIR99021 | Selleckchem | S2924 | DMSO |

| **Supplementary Table. 3. Media and reagents used in the study** | | |
| --- | --- | --- |
| **Media/Reagent** | **Product** | **Catalog no.** |
| DMEM/F12 + Glutamine + Hepes | Corning | 10-092-cm |
| 1 x Penn/Strep | Life Technologies | 10378-016 |
| 1 x Glutamine MAX | Life Technologies | 35050-061 |
| NEAA | Life Technologies | 11140-050 |
| 2ME | Life Technologies | 21985-023 |
| N2 supplement | Life Technologies | 17502-048 |
| B27 Supplement without Vitamin A | Life Technologies | 12587-010 |
| Neurolbasal | Life Technologies | 21103049 |
| TrypLE | Life Technologies | 12605-010 |

| **Supplementary Table. 4. Primer sequences and gene amplification probe sets used in our study** | |
| --- | --- |
| **Gene** | **qRT-PCR Primer sequence (5'-3')** |
| *LHX2* | F-ATCGACGAGATGGACCGCAG |
|  | R-GGTAGTAGCGGTCCGAGATC |
|  | Probe-ATGCCGTCCATCAGCAGTGACC |
| *RAX* | F-GCAAGGTCAACCTACCAGAG |
|  | R-CAGCTTCATGGAGGACACTT |
|  | Probe-CACTTAGCCCGTCGGTTCTGGAAC |
| *SIX3* | F-CACTCCCACACAAGTAGGC |
|  | R-GTGCTGGAGCCTGTTCTT |
|  | Probe-AACTGGTTTAAGAACCGGCGGCAG |
| *PAX6* | F-GTG TCT ACC AAC CAA TTC CAC AAC |
|  | R-CCC AAC ATG GAG CCA GAT G |
|  | Probe-CCACACCGGTTTCCTCCTTCACAT |
| *MITF* | F-CCTGTATGCAGATGGATGATGT |
|  | R-CCGAGACAGGCAACGTATTT |
|  | Probe-TCTTGGGCTTGATGGATCCTGCTT |
| **Gene** | **Cat no. of qRT-PCR predesigned probes from IDT** |
| *ATOH7* | Hs.PT.58.149639.g |
| *CDH6* | Hs.PT.58.40950652 |
| *FSTL4* | Hs.PT.58.3633481 |
| *SPP1* | Hs.PT.58.19252426 |
| *ISL1* | Hs.PT.58.2143768 |
| *CARTPT* | Hs.PT.58.19814336.g |
| *SOX11* | Hs.PT.58.24529615.g |
| *SNCG* | Hs.PT.58.20259255 |
| *MMP17* | Hs.PT.58.147695 |
| *GLi3* | Hs.PT.58.25033140 |
| *GLi1* | Hs.PT.58.26486279 |
| *CRX* | Hs.PT.58.40863524 |
| *CHX10 (VSX2)* | hs.pt.58.15104194 |
| *B2M* | Hs.PT.58v.18759587 |
| *CALB2* | Hs.PT.56a.26789839.g |
| *RCVRN* | Hs.PT.58.19903304 |
| *BRN3a* | Hs.PT.58.15720815 |

**Supplementary Table 5**: Differential gene expression among five conditions. **1)** L (BMP inhibitor) only, **2)** X (Wnt inhibitor) only, **3)** LSB (BMP + TGFb inhibitors), **4)** LSBX (BMP + TGFb + Wnt) inhibitors), and **5)** LSB-CHIR (BMP + TGFb inhibitors+ Wnt agonist) were evaluated in RPC cultures on day 23

| **Detector Name** | **Conditions** | **ΔΔCt** | ***p* value** | **Fold change** |
| --- | --- | --- | --- | --- |
| *ATOH7* | L vs LSB | 0.962 | 0.012 | 0.513 |
|  | L vs LSBX | 0.409 | 0.185 | 0.753 |
|  | L vs LSB-CHIR | 0.435 | 0.008 | 0.740 |
|  | X vs LSB | 0.308 | 0.094 | 0.808 |
|  | X vs LSBX | -0.246 | 0.352 | 1.186 |
|  | X vs LSB-CHIR | -0.220 | 0.034 | 1.165 |
|  | LSB vs LSBX | -0.553 | 0.133 | 1.467 |
|  | LSB vs LSB-CHIR | -0.527 | 0.034 | 1.441 |
|  | LSBX vs LSB-CHIR | 0.026 | 0.910 | 0.982 |
| *BRN3a* | L vs LSB | -0.417 | 0.067 | 1.335 |
|  | L vs LSBX | -1.471 | 0.003 | 2.772 |
|  | L vs LSB-CHIR | -1.866 | 0.008 | 3.645 |
|  | X vs LSB | 0.026 | 0.888 | 0.982 |
|  | X vs LSBX | -1.029 | 0.018 | 2.040 |
|  | X vs LSB-CHIR | -1.423 | 0.016 | 2.682 |
|  | LSB vs LSBX | -1.054 | 0.010 | 2.077 |
|  | LSB vs LSB-CHIR | -1.449 | 0.016 | 2.730 |
|  | LSBX vs LSB-CHIR | -0.395 | 0.141 | 1.315 |
| *CALB2* | L vs LSB | -0.849 | 0.018 | 1.801 |
|  | L vs LSBX | -1.554 | 0.000 | 2.937 |
|  | L vs LSB-CHIR | -1.477 | 0.002 | 2.783 |
|  | X vs LSB | 1.000 | 0.027 | 0.500 |
|  | X vs LSBX | 0.295 | 0.149 | 0.815 |
|  | X vs LSB-CHIR | 0.372 | 0.036 | 0.773 |
|  | LSB vs LSBX | -0.705 | 0.025 | 1.630 |
|  | LSB vs LSB-CHIR | -0.628 | 0.036 | 1.545 |
|  | LSBX vs LSB-CHIR | 0.078 | 0.304 | 0.948 |
| *CARTPT* | L vs LSB | 0.860 | 0.006 | 0.551 |
|  | L vs LSBX | 0.733 | 0.010 | 0.602 |
|  | L vs LSB-CHIR | -0.864 | 0.005 | 1.820 |
|  | X vs LSB | 0.942 | 0.040 | 0.520 |
|  | X vs LSBX | 0.815 | 0.053 | 0.568 |
|  | X vs LSB-CHIR | -0.782 | 0.000 | 1.719 |
|  | LSB vs LSBX | -0.127 | 0.075 | 1.092 |
|  | LSB vs LSB-CHIR | -1.724 | 0.000 | 3.303 |
|  | LSBX vs LSB-CHIR | -1.597 | 0.000 | 3.025 |
| *CDH6* | L vs LSB | -0.215 | 0.371 | 1.160 |
|  | L vs LSBX | -0.672 | 0.090 | 1.593 |
|  | L vs LSB-CHIR | -1.911 | 0.012 | 3.761 |
|  | X vs LSB | 0.049 | 0.859 | 0.967 |
|  | X vs LSBX | -0.408 | 0.264 | 1.327 |
|  | X vs LSB-CHIR | -1.648 | 0.004 | 3.133 |
|  | LSB vs LSBX | -0.457 | 0.064 | 1.373 |
|  | LSB vs LSB-CHIR | -1.697 | 0.004 | 3.241 |
|  | LSBX vs LSB-CHIR | -1.240 | 0.015 | 2.361 |
| *CRX* | L vs LSB | -2.150 | 0.003 | 4.437 |
|  | L vs LSBX | -1.928 | 0.020 | 3.806 |
|  | L vs LSB-CHIR | -2.775 | 0.005 | 6.843 |
|  | X vs LSB | -2.550 | 0.002 | 5.855 |
|  | X vs LSBX | -2.329 | 0.014 | 5.023 |
|  | X vs LSB-CHIR | -3.175 | 0.081 | 9.031 |
|  | LSB vs LSBX | 0.221 | 0.501 | 0.858 |
|  | LSB vs LSB-CHIR | -0.625 | 0.081 | 1.542 |
|  | LSBX vs LSB-CHIR | -0.846 | 0.118 | 1.798 |
| *FSTL4* | L vs LSB | -0.591 | 0.011 | 1.506 |
|  | L vs LSBX | -0.665 | 0.012 | 1.585 |
|  | L vs LSB-CHIR | -1.344 | 0.024 | 2.539 |
|  | X vs LSB | 0.642 | 0.081 | 0.641 |
|  | X vs LSBX | 0.568 | 0.104 | 0.675 |
|  | X vs LSB-CHIR | -0.112 | 0.074 | 1.081 |
|  | LSB vs LSBX | -0.074 | 0.491 | 1.053 |
|  | LSB vs LSB-CHIR | -0.754 | 0.074 | 1.686 |
|  | LSBX vs LSB-CHIR | -0.680 | 0.091 | 1.602 |
| *Gli1* | L vs LSB | -1.790 | 0.006 | 3.459 |
|  | L vs LSBX | -1.195 | 0.002 | 2.289 |
|  | L vs LSB-CHIR | -2.035 | 0.002 | 4.098 |
|  | X vs LSB | -0.345 | 0.139 | 1.270 |
|  | X vs LSBX | 0.250 | 0.069 | 0.841 |
|  | X vs LSB-CHIR | -0.590 | 0.278 | 1.505 |
|  | LSB vs LSBX | 0.595 | 0.054 | 0.662 |
|  | LSB vs LSB-CHIR | -0.245 | 0.278 | 1.185 |
|  | LSBX vs LSB-CHIR | -0.840 | 0.016 | 1.790 |
| *Gli3* | L vs LSB | -0.597 | 0.004 | 1.513 |
|  | L vs LSBX | -0.980 | 0.025 | 1.972 |
|  | L vs LSB-CHIR | -1.562 | 0.005 | 2.953 |
|  | X vs LSB | 0.118 | 0.234 | 0.922 |
|  | X vs LSBX | -0.265 | 0.255 | 1.201 |
|  | X vs LSB-CHIR | -0.847 | 0.014 | 1.799 |
|  | LSB vs LSBX | -0.383 | 0.136 | 1.304 |
|  | LSB vs LSB-CHIR | -0.965 | 0.014 | 1.952 |
|  | LSBX vs LSB-CHIR | -0.582 | 0.094 | 1.497 |
| *ISL1* | L vs LSB | -0.095 | 0.417 | 1.068 |
|  | L vs LSBX | -0.264 | 0.063 | 1.201 |
|  | L vs LSB-CHIR | -1.012 | 0.002 | 2.016 |
|  | X vs LSB | -0.426 | 0.065 | 1.343 |
|  | X vs LSBX | -0.595 | 0.025 | 1.511 |
|  | X vs LSB-CHIR | -1.343 | 0.008 | 2.537 |
|  | LSB vs LSBX | -0.169 | 0.225 | 1.125 |
|  | LSB vs LSB-CHIR | -0.917 | 0.008 | 1.888 |
|  | LSBX vs LSB-CHIR | -0.748 | 0.005 | 1.679 |
| *LHX2* | L vs LSB | -2.009 | 0.009 | 4.025 |
|  | L vs LSBX | -0.911 | 0.033 | 1.880 |
|  | L vs LSB-CHIR | -1.243 | 0.018 | 2.368 |
|  | X vs LSB | -1.826 | 0.024 | 3.546 |
|  | X vs LSBX | -0.728 | 0.118 | 1.657 |
|  | X vs LSB-CHIR | -1.061 | 0.016 | 2.086 |
|  | LSB vs LSBX | 1.098 | 0.008 | 0.467 |
|  | LSB vs LSB-CHIR | 0.765 | 0.016 | 0.588 |
|  | LSBX vs LSB-CHIR | -0.333 | 0.022 | 1.259 |
| *MMP17* | L vs LSB | -1.903 | 0.013 | 3.741 |
|  | L vs LSBX | -1.883 | 0.014 | 3.687 |
|  | L vs LSB-CHIR | -2.266 | 0.009 | 4.811 |
|  | X vs LSB | -1.373 | 0.025 | 2.590 |
|  | X vs LSBX | -1.352 | 0.027 | 2.553 |
|  | X vs LSB-CHIR | -1.736 | 0.043 | 3.331 |
|  | LSB vs LSBX | 0.021 | 0.842 | 0.986 |
|  | LSB vs LSB-CHIR | -0.363 | 0.043 | 1.286 |
|  | LSBX vs LSB-CHIR | -0.384 | 0.054 | 1.305 |
| *RCVRN* | L vs LSB | -0.764 | 0.223 | 1.698 |
|  | L vs LSBX | -2.338 | 0.017 | 5.055 |
|  | L vs LSB-CHIR | -4.277 | 0.041 | 19.388 |
|  | X vs LSB | -1.798 | 0.079 | 3.477 |
|  | X vs LSBX | -3.372 | 0.016 | 10.351 |
|  | X vs LSB-CHIR | -5.311 | 0.072 | 39.704 |
|  | LSB vs LSBX | -1.574 | 0.097 | 2.977 |
|  | LSB vs LSB-CHIR | -3.513 | 0.072 | 11.419 |
|  | LSBX vs LSB-CHIR | -1.939 | 0.177 | 3.836 |
| *SIX3* | L vs LSB | -0.163 | 0.079 | 1.120 |
|  | L vs LSBX | -0.639 | 0.006 | 1.557 |
|  | L vs LSB-CHIR | -0.990 | 0.003 | 1.986 |
|  | X vs LSB | 0.215 | 0.243 | 0.862 |
|  | X vs LSBX | -0.260 | 0.184 | 1.198 |
|  | X vs LSB-CHIR | -0.612 | 0.001 | 1.528 |
|  | LSB vs LSBX | -0.475 | 0.001 | 1.390 |
|  | LSB vs LSB-CHIR | -0.827 | 0.001 | 1.774 |
|  | LSBX vs LSB-CHIR | -0.351 | 0.004 | 1.276 |
| *SNCG* | L vs LSB | -0.351 | 0.235 | 1.276 |
|  | L vs LSBX | -0.354 | 0.036 | 1.278 |
|  | L vs LSB-CHIR | -0.411 | 0.025 | 1.330 |
|  | X vs LSB | 0.102 | 0.737 | 0.932 |
|  | X vs LSBX | 0.099 | 0.629 | 0.933 |
|  | X vs LSB-CHIR | 0.042 | 0.794 | 0.971 |
|  | LSB vs LSBX | -0.003 | 0.991 | 1.002 |
|  | LSB vs LSB-CHIR | -0.060 | 0.794 | 1.042 |
|  | LSBX vs LSB-CHIR | -0.057 | 0.212 | 1.040 |
| *Sox11* | L vs LSB | -2.023 | 0.005 | 4.063 |
|  | L vs LSBX | -1.797 | 0.003 | 3.474 |
|  | L vs LSB-CHIR | -1.833 | 0.013 | 3.564 |
|  | X vs LSB | -2.054 | 0.002 | 4.152 |
|  | X vs LSBX | -1.828 | 0.000 | 3.550 |
|  | X vs LSB-CHIR | -1.865 | 0.458 | 3.642 |
|  | LSB vs LSBX | 0.226 | 0.157 | 0.855 |
|  | LSB vs LSB-CHIR | 0.189 | 0.458 | 0.877 |
|  | LSBX vs LSB-CHIR | -0.037 | 0.861 | 1.026 |
| *SPP1* | L vs LSB | -0.986 | 0.094 | 1.981 |
|  | L vs LSBX | 0.086 | 0.816 | 0.942 |
|  | L vs LSB-CHIR | -1.270 | 0.076 | 2.412 |
|  | X vs LSB | -1.127 | 0.007 | 2.183 |
|  | X vs LSBX | -0.054 | 0.628 | 1.038 |
|  | X vs LSB-CHIR | -1.411 | 0.255 | 2.659 |
|  | LSB vs LSBX | 1.072 | 0.001 | 0.475 |
|  | LSB vs LSB-CHIR | -0.284 | 0.255 | 1.218 |
|  | LSBX vs LSB-CHIR | -1.357 | 0.017 | 2.561 |
| *RAX* | L vs LSB | -3.264 | 0.000 | 9.607 |
|  | L vs LSBX | -1.326 | 0.027 | 2.507 |
|  | L vs LSB-CHIR | -1.526 | 0.002 | 2.880 |
|  | X vs LSB | -2.416 | 0.000 | 5.336 |
|  | X vs LSBX | -0.478 | 0.167 | 1.392 |
|  | X vs LSB-CHIR | -0.678 | 0.002 | 1.600 |
|  | LSB vs LSBX | 1.938 | 0.013 | 0.261 |
|  | LSB vs LSB-CHIR | 1.738 | 0.002 | 0.300 |
|  | LSBX vs LSB-CHIR | -0.200 | 0.473 | 1.149 |
| *MITF* | L vs LSB | -1.349 | 0.001 | 2.548 |
|  | L vs LSBX | -1.335 | 0.002 | 2.523 |
|  | L vs LSB-CHIR | -1.567 | 0.002 | 2.963 |
|  | X vs LSB | -0.355 | 0.014 | 1.279 |
|  | X vs LSBX | -0.340 | 0.018 | 1.266 |
|  | X vs LSB-CHIR | -0.572 | 0.107 | 1.487 |
|  | LSB vs LSBX | 0.014 | 0.837 | 0.990 |
|  | LSB vs LSB-CHIR | -0.218 | 0.107 | 1.163 |
|  | LSBX vs LSB-CHIR | -0.232 | 0.099 | 1.175 |
| *PAX6* | L vs LSB | -1.015 | 0.003 | 2.021 |
|  | L vs LSBX | -0.960 | 0.003 | 1.945 |
|  | L vs LSB-CHIR | -1.303 | 0.010 | 2.467 |
|  | X vs LSB | -0.291 | 0.028 | 1.224 |
|  | X vs LSBX | -0.236 | 0.031 | 1.178 |
|  | X vs LSB-CHIR | -0.579 | 0.174 | 1.493 |
|  | LSB vs LSBX | 0.055 | 0.488 | 0.962 |
|  | LSB vs LSB-CHIR | -0.287 | 0.174 | 1.220 |
|  | LSBX vs LSB-CHIR | -0.343 | 0.129 | 1.268 |

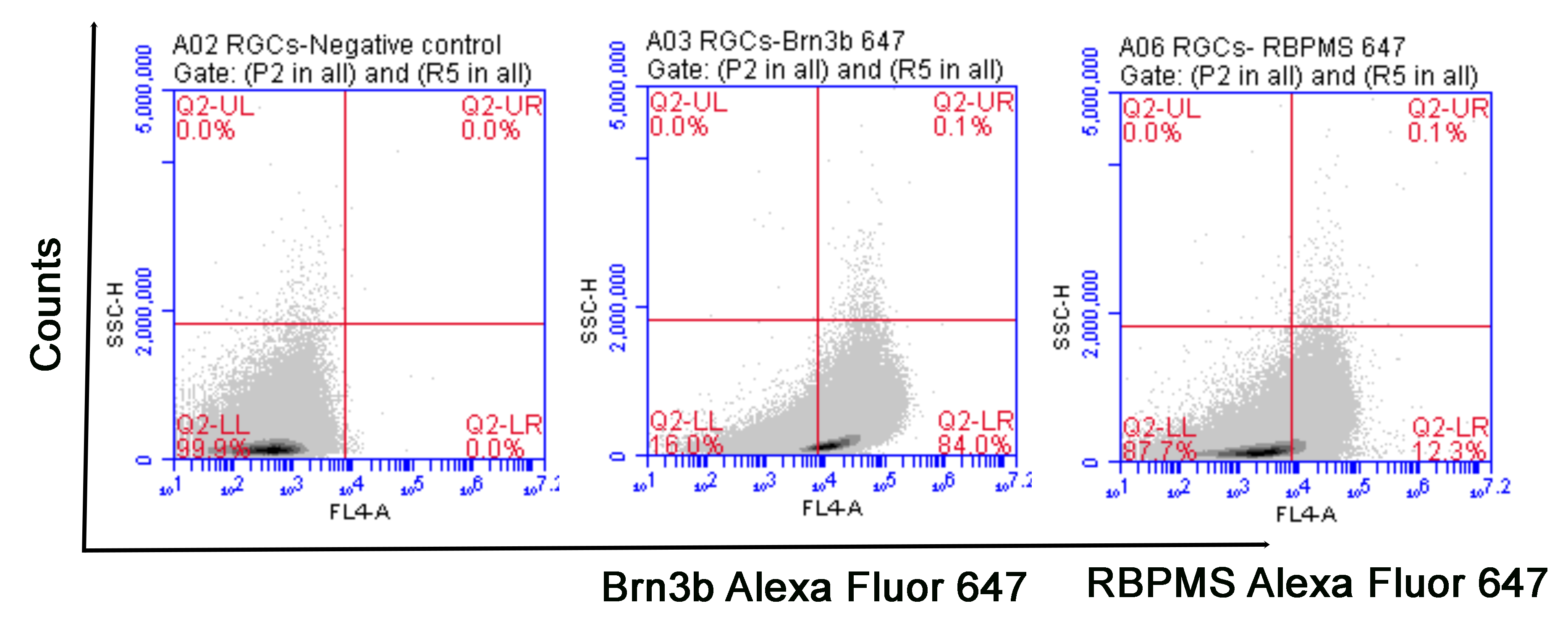

**Supplementary Fig. 1.** Flow cytometry analysis showing percentage of Brn3b positive and RBPMS positive iPSC-RGCs.

**Additional information: Protocol for differentiation of iPSC-RGCs using small molecules and proteins**

**Protocol time line:**

D-3 to-1	Day 0-15	Day 16-21	Day 22-23	Day 24	Day 25-26	Day 27-36

RPC conditions	RGC differentiation

Initiation	RPC induction and differentiation	RGC initiation	cross-hatching	RGC differentiation	RGC maturation

**iPSC initiation: day-3 to day-1**

Day1: First MEFs cell depletion**:** When iPSC cells reach 80% confluent, apply 1ml of TrypLE solution and incubate at 37^0^C for 4-5 minutes. Remove TrypLE medium after incubation and replate iPSCs into a 100mm dish precoated with matrigel, in medium containing: 80% HES / 20% MEF-CM + 20ng/ml bFGF + 4uM Y27632. Incubate the plate in 37^0^C, 5%CO_2_ and 5%O_2_ incubator.

Day2: Second MEFs cell depletion**:** iPSC cells should be over 80% confluent in this step. 2ml of TrypLE solution was added to 100mm dish and incubated at 37^0^C for 4-5 minutes. After TrypLE treatment, iPSC cells were replated into one well of a 6 well plate, precoated with Matrigel (seeding density of 1.5X10^6^ cells per well), in medium containing: 80%HES / 20% MEF-CM + 20ng/ml bFGF + 5ng/ml stable bFGF + 4uM Y27632 and incubated in 37^0^C, 5%CO_2_ and 5%O_2_ incubator.

Day3: The spent medium is replaced with a fresh medium without Y27632. (80%HES / 20% MEF-CM + 20ng/ml bFGF + 5ng/ml stable bFGF). The plate is moved from the 5% O_2_ incubator to the regular O_2_ incubator maintained at 37^0^C, 5%CO_2_ incubator (the night before) to continue with RPC induction and differentiation.

**1. RPC Induction and Differentiation: day 0-21**

**A. Day0-d3**:

Change medium daily with RPC induction media containing: 0.1µM LDN, 10µM SB, 2µM XAV, 10mM NIC, 10ng/ml IGF1. Add all small molecule and protein into each day’s medium aliquot freshly (just before medium change).

**RPC induction media:**

|  | **Reagent** | **Final Concentration** |
| --- | --- | --- |
| 1 | Regular DMEM/F12 |  |
| 2 | 1 x Penn/Strep (P/S; 100X) | 1% |
| 3 | 1 x Glutamine MAX (100X) | 1% |
| 4 | NEAA (MEM Non Essential Amino Acid) | 1% |
| 5 | 2-Mercaptoethanol (55mM stock) | 0.1mM |
| 6 | N2 supplement (1%) | 1% |
| 7 | B27 Supplement without Vitamin A (2%) | 2% |

**Day 0-3: medium components**

|  | **Component** | **Final Concentration** |
| --- | --- | --- |
| 1 | RPC induction media | |
| 2 | LDN | 0.1µM |
| 3 | SB | 10µM |
| 4 | XAV939 | 2µM |
| 5 | Nicotinamide (NIC) | 10mM |
| 6 | IGF1 | 10ng/ml |

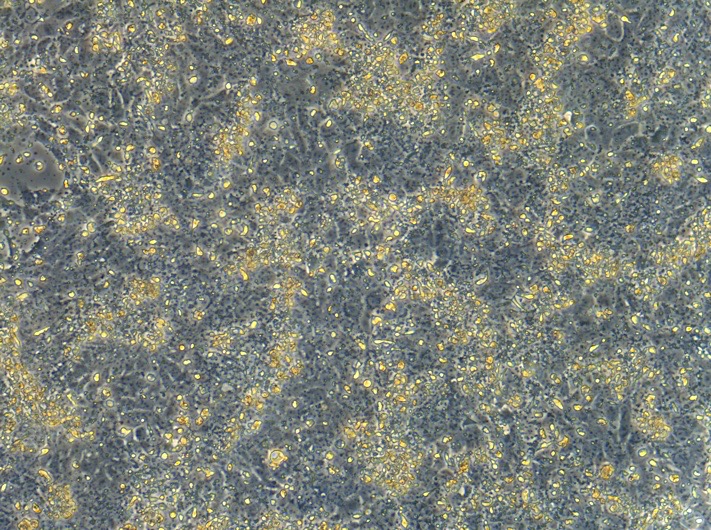

*By Day3, cells continued to expand, with cells reaching more than 100% confluence* (Mag- 10X)*.*

**B. Day 4-21:**

Change medium daily with RPC induction media containing: 0.1µM LDN, 10µM SB, 2µM XAV, 1.5µM

CHIR, 10ng/ml IGF1, and 10ng/ml bFGF. Add all small molecule and protein into each day’s medium aliquot freshly (just before medium change).

**Day 4-21: media components**

|  |  | **Final Concentration** |
| --- | --- | --- |
| 1 | RPC induction media | |
| 2 | LDN | 0.1µM |
| 3 | SB | 10µM |
| 4 | XAV939 | 2µM |
| 5 | CHIR99021 | 1.5µM |
| 6 | IGF1 | 10ng/ml |
| 7 | bFGF | 10ng/ml |

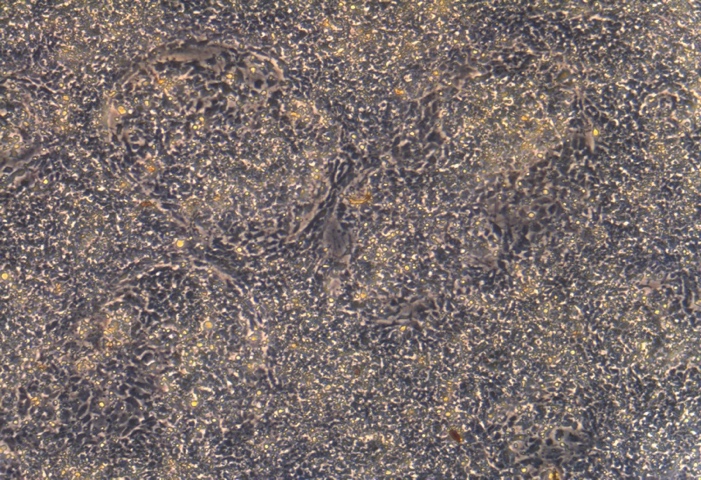

*By Day19, more rosette like cell clusters formed (starting around D15)*, (Mag- 10X)

**2. RGC initiation and differentiation Day22-36**

**A. Day22-23: RGC initiation**

Change medium daily with RGC induction media containing: 3μM DAPT, 250ng/ml Shh, and 100ng/ml FGF8.

**RGC induction media**

|  | **Reagent** |  |
| --- | --- | --- |
| 1 | Regular DMEM/F12 | 50% |
| 2 | Neuraobasel media | 50% |
| 3 | 1 x Penn/Strep (P/S; 100X) | 1% |
| 4 | 1 x Glutamine MAX (100X) | 1% |
| 7 | B27 Supplement without Vitamin A (2%) | 2% |

**Day22-23: medium Components**

|  | **Component** | **Final Concentration** |
| --- | --- | --- |
| 1 | RGC induction media | |
| 2 | DAPT | 3μM |
| 3 | Shh | 250ng/ml |
| 4 | FGF8 | 100ng/ml |

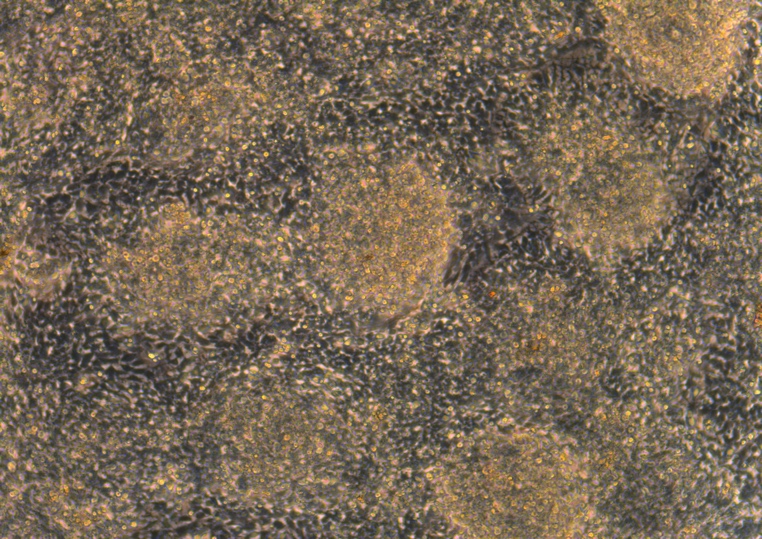

*Day24 before crosshatching, abundant cell clusters with crowded packed cells forming multi layers (light yellow parts).*

**B. D24 RGC Differentiation: cross-hatching and replat**

On the Day 24

1. On ice, add 30ml of Leibovitz’s (LM) medium L15 + 1M Glucose into 50ml tube.

2. Remove 6 well cell plate from incubator, wash the well once with cold LM + Gluc. (LMG) media (2mL/well).

3. Add 2 mL of LMG media to each well of 6-well dish and scrape using a p1000 tip with the pipette, cross the bottom firmly, cross horizontal lines parallelly and closely to the full bottom surface, then cross again vertically, to dislodge cells into small clusters.  Lift dish up to ensure all cells are being removed (>90%). Draw medium up and blew down 3 times, each time to rains off cells from different bottom area, then do two more times to break cells down to even and small clusters. Check under the scope, if cell cluster still big, blew up/down cells two more times.

4. remove all cells with LM+Gluc from above into 20mL of LMG media in a 50ml tube, and spin down at 3000rpm for 3min.

5. remove supernatant and resuspend cells pellets in 8ml Day24 medium + 4uM Y27632), blew cells up and down about 4-6 times. Add enough Day24 medium + Y27632 for plating. Cell seeding density: about 1X10^5^/well of 6 well plates.

**Day24: medium Components**

|  | **Component** | **Final Concentration** |
| --- | --- | --- |
| 1 | RGC induction media | |
| 2 | DAPT | 3 μM |
| 3 | Cyclopamine | 0.5uM |
| 3 | Follistatin | 100ng/ml |

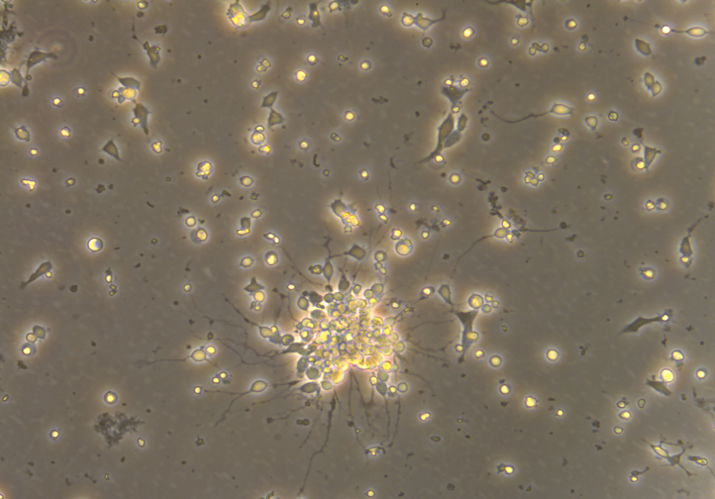

*Day24, about 3 hours after crosshatching, neuron axon cell bodies start to form* (Mag- 20X)

**C. RGC Differentiation, Day25-26**

Change medium daily with RGC induction media containing: 3μM DAPT and 100ng/ml Follistatin

**Day25-26: medium components**

|  | **Component** | **Final Concentration** |
| --- | --- | --- |
| 1 | RGC induction media | |
| 2 | DAPT | 3 μM |
| 3 | Follistatin | 100 ng/ml |

**3. RGC Maturation, Day27-36**

Change medium every 2-3 days with RGC induction media containing: 3μM DAPT, 10μM Y27632, 400μM cAMP, 5μM Forskolin, 40ng/ml BDNF, 5ng/ml NT4 and 10ng/ml CNTF

**Day27-36: medium components**

|  | **Component** | **Final Concentration** |
| --- | --- | --- |
| 1 | RGC induction media | |
| 2 | DAPT | 3μM |
| 3 | Y27632 | 10μM |
| 4 | cAMP | 400μM |
| 5 | Forskolin | 5μM |
| 6 | BDNF | 40ng/ml |
| 7 | NT4 | 5ng/ml |
| 8 | CNTF | 10ng/ml |

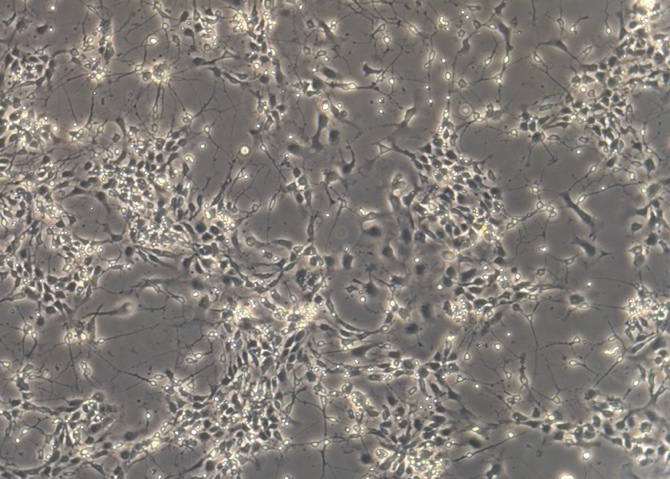

Day27, numerous ganglion cell bodies with extended axons (Mag- 10X).

**4. Mature RGC maintenance (from Day37)**

Change medium twice a week with RGC induction media containing: 3μM DAPT, 10μM Y27632

**From D37: medium components**

|  | **Component** | **Final Concentration** |
| --- | --- | --- |
| 1 | RGC media | |
| 2 | DAPT | 3 μM |
| 3 | Y27632 | 10 μM |

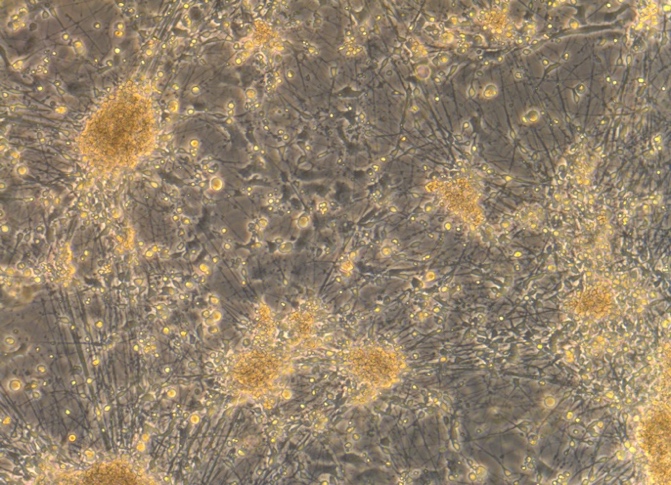

*Day37, matured RGC with long axons, RGC cell bodies migrate forming clusters* (Mag- 10X)*.*
